# Modeling the microbiome of Utah’s Great Salt Lake: A regression analysis of key abiotic factors impacting growth of *Dunaliella* green algae in the GSL’s South Arm

**DOI:** 10.1101/2024.02.10.579688

**Authors:** Catherine G. Fontana, Vanessa G. Maybruck, Rachel M. Billings, Cresten B. Mansfeldt, Elizabeth J. Trower

## Abstract

Since the mid-1800s, Utah’s Great Salt Lake (GSL) has undergone dramatic changes. Due to the effects of climate change and an increase in agricultural, industrial, and residential water usage to support population growth, the present water level has fallen to about one-fourth of its highest recorded level in 1987 [1, 2]. As Earth’s global air and water temperatures continue to rise, evaporation rates from this closed basin will also rise, thus increasing the salinity of this already hypersaline lake. A shift in water chemistry from its current salinity of 15% to a halite saturation of 30% will negatively impact the populations of *Dunaliella viridis* – a halophilic species of green algae that form the basis of the simple but delicate food web in the South Arm of the GSL. Disruption of the *D. viridis* population through increased water temperature and salinity will spur a negative cascade throughout the food chain by reducing brine shrimp populations and thereby threaten local and migratory bird populations. Since increasing water temperature and salinity can have such deleterious ramifications on both *D. viridis* and the overall lake ecosystem, a predictive model that maps the impact of changing water temperature and salinity to specific growth values for *D. viridis* is needed for forecast-assisted management. In support of this goal, we developed a multiple linear regression model using twelve years of observational data consisting of chlorophyte (of which *Dunaliella* are the dominant species) population concentrations under co-varying water temperature and salinity. The resulting fitted data produced an *R*^2^ value of 0.17 with a RMSPE of 100.704, and additional diagnostics were conducted to verify the model. Overall, this model predicts that chlorophyte populations will decrease by 0.41 *μ*g/L for each 1% increase in salinity and decrease by 0.74 *μ*g/L for each 1°C increase in water temperature up to the extinction point of 30% salinity and 45°C. One limitation of the linear regression model is its inability to capture trace algal population concentrations at 0 *μ*g/L. To address this, we also developed a zero-inflated Poisson regression model, which predicts similar decreases in chlorophyte populations for increasing water temperature and salinity as the linear regression model. The fitted data for this model produced a pseudo-*R*^2^ value of 0.35 with a RMSPE of 90.026. This model predicts that chlorophyte populations will decrease by 0.16 *μ*g/L for each 1% increase in salinity and decrease by 0.13 *μ*g/L for each 1°C increase in water temperature up to the extinction point of 30% salinity and 45°C. Even for a limited climate change scenario of an increase in air/water temperature of 2.5°C and an associated increase in salinity by 7.5%, the linear regression model predicts a potential loss of ∼224, 000 kg total of chlorophytes from the South Arm of the GSL (based on the median chlorophyte concentration between 2001 and 2006), while the Poisson regression model predicts a potential loss of ∼173, 200 kg of chlorophytes. Continued research will include model selection and error quantification. More broadly, future work aims to constrain chlorophyta population predictions based on *D. viridis* growth limits under maximum water temperature and salinity thresholds obtained from controlled laboratory experiments, which can be used to identify a microbial tipping point of the GSL.

## 1 Introduction

The Great Salt Lake (GSL) sits in the historic Lake Bonneville Basin in northwest Utah. During the end of the Last Glacial Maximum (30,000 to 13,000 years ago), Lake Bonneville was an expansive paleolake that covered the entire northwestern corner of present-day Utah and extended into northeast Nevada and southeast Idaho. Today, three smaller lakes remind us of the expanse of what once was: Sevier Lake, Utah Lake, and the Great Salt Lake, the latter of which being the largest (2, 100km^2^ as of 2021) [3]. These lakes and their associated network of rivers and streams comprise the modern Lake Bonneville Basin, which is the largest contiguous watershed in North America [4].

The Great Salt Lake is the largest lake in the western United States and the second saltiest lake in the world [4]. Its high salinity is a result of it being a terminal lake: it receives input from freshwater streams (namely the Jordan, Weber, and Bear Rivers) but does not have any outflow [5, 6, 7]. Since its formation, minerals have been weathered from nearby mountains have eroded, delivering dissolved ions such as Na and Mg into the GSL. On-going evaporation from this closed basin has systematically concentrated these dissolved ions within the lake’s water, bringing it to its present characteristic salinity.

The GSL is a major economic driver in Utah, contributing approximately 2.5 billion dollars to the state’s economy annually [8]. In addition to being a top tourist destination in the state of Utah, the lake is also harvested for a variety of resources used in different industries including salt (roads), magnesium chloride (steel), lithium (batteries), potassium sulfate (fertilizer), and brine shrimp (commercial fisheries) [9]. The GSL’s impacts also extend beyond its immediate vicinity: it is the source of up to ten percent of snow deposited at Utah’s ski resorts [9]. The GSL is also ecologically important and is considered to be the most important migratory bird stop in North America [4, 10]. Each year, approximately 10 million birds of 250 species stop at the GSL to feed on its abundant brine fly and brine shrimp populations as part of their migration [4, 10].

In 1959, a causeway was constructed across the lake, bisecting it into what is now known as the North Arm and South Arm (Figure 1). This causeway created an artificial barrier that resulted in distinct conditions between the GSL’s arms. The salinity of the North Arm approaches 30% (halite saturation) due to limited freshwater inflow [5, 6]. The salinity of the lake’s South Arm varies between 8 − 17% due to freshwater inflow from the Jordan, Weber, and Bear Rivers [6]. Both North Arm and South Arm ecosystems are two to seven times saltier than the ocean (3% salinity) and host similar halophilic species. However, research has suggested that the North Arm’s food web is less functionally redundant (fewer species, as related to those on South Arm) than the South Arm’s resulting from selection pressures created by its near-saturation levels of salinity [4].

**Figure 1:**
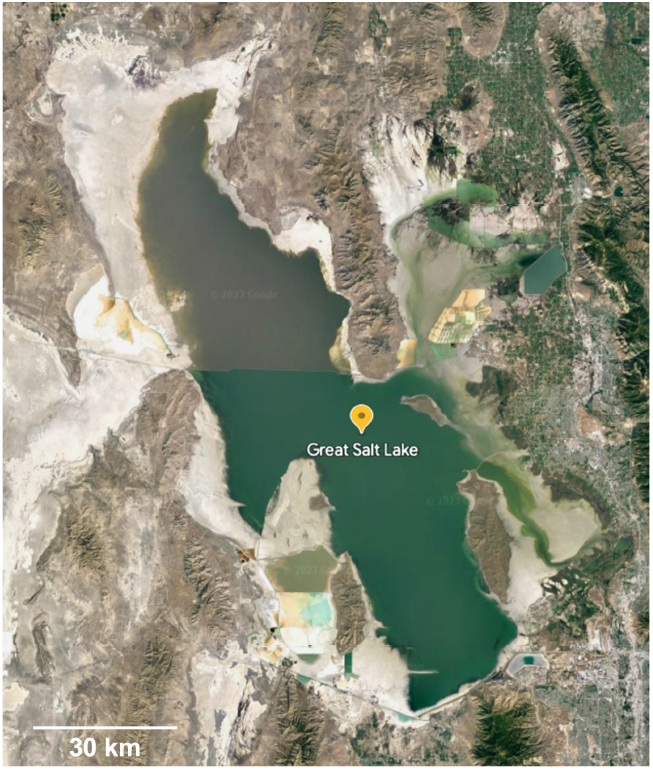
Aerial photo of the GSL in 2023 that shows clear delineation between the North Arm and South Arm salinity gradient created by the causeway erected in 1959. Satellite image obtained via Google Earth.

### 1.1 An Ecosystem In Peril

As a terminal lake, the GSL is highly susceptible to climate change and water diversion [11]. Damming and water diversion of feeder freshwater streams began in the late 1800s to meet industrial, agricultural, and municipal demands [8, 11]. Currently, more than two-thirds of the GSL’s incoming freshwater streamflow is diverted with 74% used for agriculture [8]. Since the summer months of 1985, the lake elevation has dropped by 20 feet and has fallen to about one-fourth of its highest recorded value [1, 2]. At its current loss rate, the GSL is estimated to dry up in as few as 5 years [8]. The loss of this unique ecosystem would devastate both the economy and environment of Salt Lake City and greater Utah.

### 1.2 Microbiology of the Great Salt Lake

Although it has been called ‘America’s Dead Sea’ [12, 13], the Great Salt Lake contains a myriad of diverse lifeforms such as brine flies, brine shrimp (*Artemia*), free-floating phytoplankton, and bottom-dwelling cyanobacteria. Given the differences in salinity, two related but separate ecosystems have formed in the North and South Arms of the GSL. The South Arm’s relatively lower salinity has allowed a more diverse ecosystem to establish that supports annual blooms of the algae *Dunaliella viridis*, a critical food source for *Artemia* [4]. A simpler, more streamlined network of organisms - anchored by a more halophilic algae, *Dunaliella salina* - has established on the North Arm due to its near-saturation salinity [4].

Nutrient cycling and decomposition are critical biochemical processes that sustain balance in the GSL and are driven by the resident microorganisms [4]. *Dunaliella* is often the sole primary producer in its environment [14], forming the foundation of many food webs and supporting other organisms. The food web of the Great Salt Lake is sustained by brine flies and brine shrimp, for which free-floating microbial algae such as the *Dunaliella* species are a key source of food. These flies and shrimp, in turn, are important sources of food for the millions of migratory birds that visit the lake each year. In addition to serving as the primary base of the food web, these microbes also contribute to nutrient turnover and decomposition in the lake, further supporting the lake’s ecosystems through helping to maintain their complex biochemical balance [4].

### 1.3 Project Aim and Significance

Despite their importance as primary producers in the GSL and other saline ecosystems, *Dunaliella* are understudied in their natural environments [14]. However, research has demonstrated that *Dunaliella* are sensitive to changes in their environment [4, 15, 16]. Thus, *Dunaliella* can be used as an indicator species to both identify minute shifts in environmental conditions and make predictions about future ecosystem function of the GSL. This research leverages existing observational data to predict how *Dunaliella* may be impacted by the ongoing effects of anthropogenic change, including water diversion and global climate change. The findings from this research may serve as part of future impact and conservation management studies, thus supporting the conservation of the GSL as a whole.

## 2 Methods

We began by identifying existing data sets from peer-reviewed literature. Papers from the initial search were filtered based on their suitability for multiple regression analysis of *D. salina* or *D. viridis* growth, specifically their inclusion of co-varying data on water temperature and salinity. The data set published by Belovksy et al. 2011 was selected because it contained 58 observational data points from 1994 to 2006 on South Arm water temperature, salinity, and chlorophyte concentrations [16]. As *Dunaliella* comprises 93.5% of the chlorophyta biovolume in the GSL South Arm [17], chlorophyte data will be used to represent *Dunaliella* populations in this analysis.

The histograms displayed in Figure 2 show the distributions of considered variables from the chosen data set. Most of the salinity values were between 8 and 9%, which is near the ideal salinity for *D. viridis*. Water temperature values ranged from around 0 to around 25°C. This data set has some associated challenges. First, data was not continuously measured over the data set’s 12 year span; additionally, the distribution of chlorophyte data was heavily skewed towards zero. It is suspected that the large number of 0 *μ*g/L measurements for chlorophyte concentration are due to instrument detection limits.

**Figure 2:**
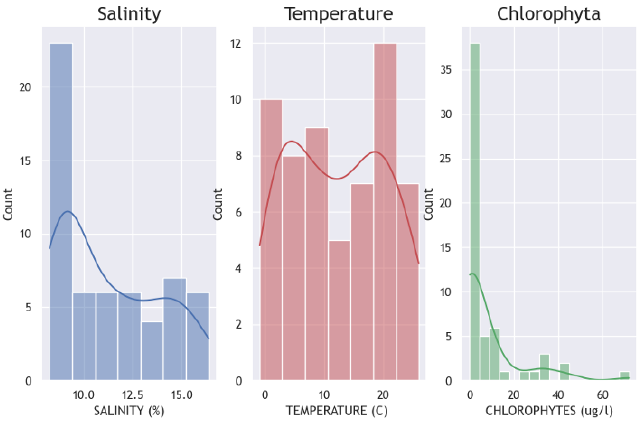
Histograms of the distributions of salinity, water temperature, and chlorophyte biomass concentration data. Chlorophyte data are heavily skewed to 0 *μ*g/L.

### 2.1 Data Processing and Modeling Methods

Since our goal is to predict *Dunaliella viridis* population levels from water temperature and salinity, we conducted our analysis using multiple regression models, which learn patterns in the data and are commonly used for prediction and explanation [18, 19]. Specifically, we developed regression models with chlorophyte concentration as the response variable and water temperature and salinity as the predictor variables. To make the data suitable for regression, we first extracted the data of interest (water temperature, salinity, and chlorophyte concentration) from the larger data set, removed the records with missing chlorophyte concentration measurements, and converted all data to a numeric data type. Since prediction is our goal and we would like to calculate the root mean squared prediction error (RMSPE) to determine the model’s prediction accuracy, we randomly split the data into two sets of approximately equal size: the set on which we trained the model (training set) and the set on which we tested the model (testing set) [20, 21]. We began by fitting a multiple linear regression model and additionally fitted a zero-inflated Poisson regression model to obtain a better model fit. Both models are described in more detail in the following sections.

## 3 Results

Here we present our regression analysis of the data set thus far, including linear and zero-inflated Poisson regression models.

### 3.1 Multiple Linear Regression Model

Using the linear_model.LinearRegression() function from Python’s scikit-learn library, we generated a multiple linear regression model with *R*^2^ = 0.17. Since the *R*^2^ value is much less than 1, we can see that the model is likely not a good fit, since a low *R*^2^ value means that much of the variability in the data is not being explained by the model. This conclusion is verified by Figures 3–4.

**Figure 3:**
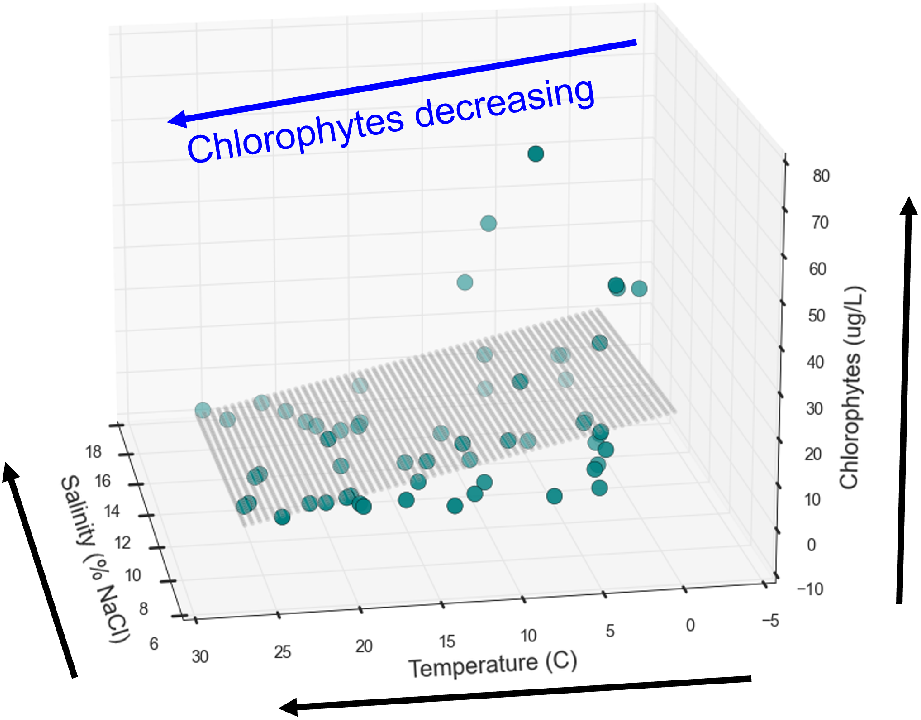
3-D rendering of the linear regression model with chlorophyte concentration as the response variable and water temperature and salinity as predictors. As water temperature and salinity increase, chlorophyte concentration decreases. *R*^2^ = 0.17.

If the fit were good, then all of the points in Figure 3 would lie on or near the plane. Instead, we see significant over-fitting and under-fitting, resulting in the large residuals that cause the poor *R*^2^ value. Furthermore, if the fit were good, then the model should predict chlorophyte concentrations from the water temperature and salinity that are close to the actual chlorophyte concentrations. In that case, we would know that the model is accurately learning informative trends within the data. Graphically, a plot of fitted chlorophyte concentrations versus actual concentrations should display a tight fit along the line *y* = *x*. However, heavy clustering close to the origin is noted (Figure 4), likely because 22 of the 58 records in the data set have chlorophyte concentrations measured as 0 *μ*g/L. We handle this problem in the zero-inflated Poisson regression model in the following section.

**Figure 4:**
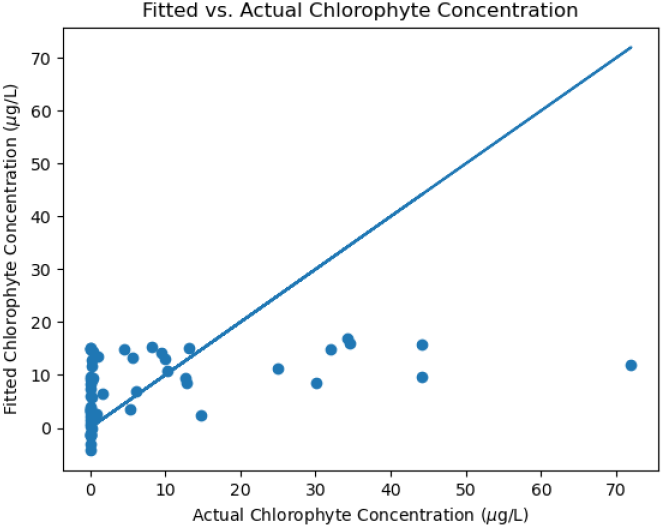
Plot of fitted chlorophyte concentrations predicted by the linear regression model vs. actual chlorophyte concentrations from the data. If the model were a good fit, then the points in this plot would be tightly grouped around the line *y* = *x*.

Despite the poor fit of the linear model, we can still learn from it. In Figure 3, we see that increasing water temperatures and salinity correspond with decreasing chlorophyte concentration. This result is confirmed, and more easily visualized, by plotting water temperature and salinity individually against chlorophyte concentration (Figure 5).

**Figure 5:**
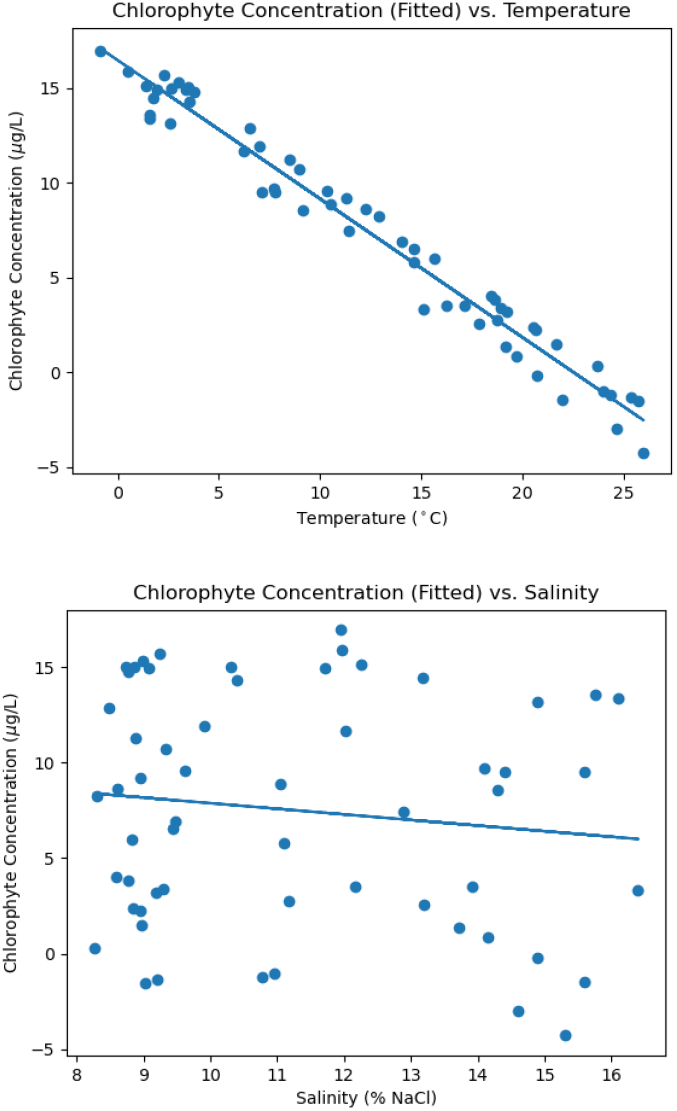
2-D renderings of the linear regression model, showing the relationships between water temperature, salinity, and chlorophyte concentration. The relationship between salinity and chlorophyte concentration appears nonlinear, whereas the relationship between water temperature and chlorophyte concentration appears linear.

From Figure 5, we can see that water temperature and chlorophyte concentration share a linear relationship, whereas salinity and chlorophyte concentration do not. Therefore, the poor fit of the linear regression model is likely due to a nonlinear relationship between salinity and chlorophyte concentration, whereas, if we were modeling the effect of water temperature alone on chlorophyte concentration, then a linear model would likely be an accurate fit. In addition to these more general conclusions about variable relationships, we can also draw more specific conclusions from our linear model. For instance, we can see that a 1°C increase in water temperature is associated with a 0.74 *μ*g/L decrease in chlorophyte concentration, while a 1% increase in salinity is associated with a 0.41 *μ*g/L decrease in chlorophyte concentration. Unsurprisingly, when we ran our *t*-tests for statistical significance, we found that the *p*-value for water temperature is statistically significant (*p* = 0.001), while the *p*-value for salinity is not (*p* = 0.559) for a significance level of *α* = 0.05. This confirms our previous observation that temperature is a good predictor for chlorophyte concentration if we use a linear regression model, while salinity is not, and indicates that temperature should be included in the linear model while salinity should be excluded.

To assess the model’s effectiveness at prediction in relation to other models, we also computed the RMSPE by running the linear regression model on the testing set and comparing the predicted chlorophyte concentration values with the actual chlorophyte concentrations in the testing set. In doing so, we found that the linear regression model has RMSPE = 100.704, a value which will be more meaningful when compared with the RMSPE in the zero-inflated Poisson regression model, which is described in the next section.

### 3.2 Zero-Inflated Poisson Regression Model

As we see in Figure 2, the chlorophyte concentrations are heavily skewed towards 0 *μ*g/L, an attribute that is not handled well by the linear regression model. Therefore, we elected to fit a second model, this time a zero-inflated Poisson regression model, which is specifically equipped to handle data sets in which the response variable has an excessive number of zero entries. We achieved this by using the ZeroInflatedPoisson() function from Python’s statsmodels library, specifically electing to use a Poisson model because of its common usage in modeling counts and rates (like we have here). On a surface level, the zeroInflated Poisson() function first runs a logistic regression model to determine the probability of the chlorophyte concentration being 0 *μ*g/L. Then it incorporates this probability into two different probability mass functions (PMFs), which are derived from the PMF for the standard Poisson regression model; one PMF is for cases where the chlorophyte concentration is 0 *μ*g/L and the other is for cases with strictly positive chlorophyte concentrations. Together, the PMFs are used to generate a two-part probability distribution that is used to train the zero-inflated Poisson model.

In Poisson regression modeling, we assume that the response variable (chlorophyte concentration in this case) is not normally distributed. Therefore, we cannot compute the *R*^2^ value as in the linear regression model and instead compute the pseudo-*R*^2^ value, which is comparable to the *R*^2^ value given by the linear regression model. Upon running the model, we see that the pseudo-*R*^2^ value is 0.35. This is still relatively low, since the ideal *R*^2^ value is 1, and demonstrates the need for continued data analysis and model fitting. However, it is much better than the *R*^2^ = 0.17 from the linear model. While it is difficult to assess whether the better model fit was a result of using a Poisson model rather than a linear model or of appropriate handling of excess zeros in the data, future model fitting should incorporate one or both of these considerations. Furthermore, we also found that both water temperature and salinity are statistically significant, with *t*-tests returning *p <<* 0.05, which indicates that both are necessary for the model.

With a better model fit established, we can trust our conclusions from the Poisson model more so than the ones we made previously from the linear regression model. The general trend between the two models is the same: increasing water temperature and/or salinity corresponds with a decrease in chlorophyte concentration. The difference between the models’ conclusions instead resides in the magnitude of these changes. Specifically, an increase of 1°C in water temperature is associated with a 0.13 *μ*g/L decrease in chlorophyte concentration, while a 1% increase in salinity is associated with a 0.16 *μ*g/L decrease in chlorophyte concentration (compared to 0.74 and 0.41 *μ*g/L, respectively, for the linear model). Keeping in mind our goal of prediction, we again computed the RMSPE for the Poisson model and found that RMSPE_Poisson_ = 90.026 *<* 100.704 = RMSPE_linear_. In other words, the Poisson model has less prediction error than the linear model and is superior in terms of prediction. With that being said, the Poisson model still exhibits a significant prediction error and requires further refinement to obtain more accurate predictions (Figure 6).

**Figure 6:**
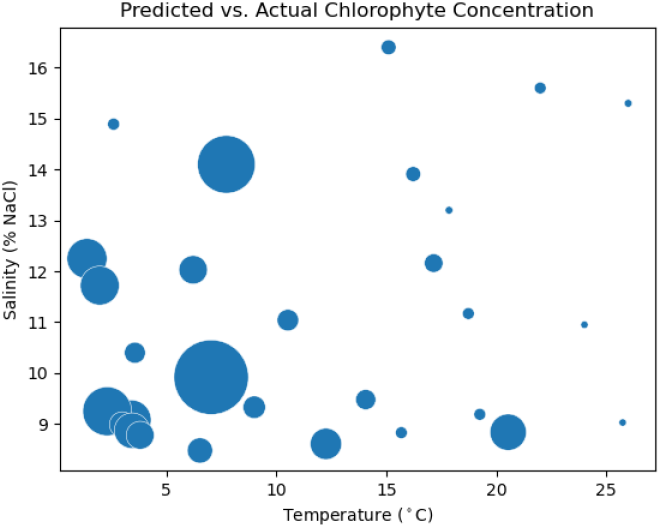
Absolute prediction error, as bubbles, plotted along the water temperature and salinity continuums. Error is computed as |actual [chlorophyte] − predicted [chlorophyte] | Larger bubbles indicate larger prediction error.

## 4 Discussion

Daily monitoring by the U.S. Geological Survey has proven paramount in tracking the GSL’s water level decline over decades, but few studies have similarly tracked microbial dynamics over time. To our knowledge, Belovsky et al. is the first long-term, in situ, observational study of various phytoplankton – including chlorophytes – in the GSL [16]. Using this existing data set, this research quantifies chlorophyte population dynamics in relation to the water temperature and salinity in the South Arm of Utah’s GSL.

Our analyses forecast that chlorophyte populations will decrease between 0.13 to 0.74 *μ*g/L with every 1°C increase in water temperature. We additionally project that chlorophyte populations will decrease between 0.16 to 0.41 *μ*g/L with every 1% increase in salinity. In this analysis, we have assumed that lake water temperature changes at the same magnitude as local air temperature. Even for a limited climate change scenario of an increase in air/water temperature of 2.5°C and an associated increase in salinity by 7.5%, these models predict a potential loss between 173, 200 kg and 224, 000 kg total of chlorophytes in the South Arm (Poisson regression and linear regression, respectively), based on the median chlorophyte concentration between 2001 and 2006.

These projected decreases in chlorophyte populations will cascade throughout the GSL’s food web. Any modulation of the chlorophyte population will disrupt brine shrimp populations, which are an essential food source for the over 10 million migratory birds that rely on the GSL for their annual stopover. In short, a loss of chlorophyte biomass could significantly and negatively impact one of the the most important migratory bird locations in North America.

### 4.1 Limitations

There are several important limitations in both data availability and analysis. Since *Dunaliella* are understudied, it was difficult to identify a data set that was suitable for multiple regression. Despite our extensive literature review, our selected data set contains data for chlorophytes, not *Dunaliella*, and we based our analysis on the assumption that *Dunaliella* comprise a sufficient percentage of chlorophytes in the GSL to substitute the two groups of microbes with each other. However, perhaps our data analysis would have yielded different results if there were more data available that are specific to *Dunaliella*. Additionally, there were many missing values in the data set. While 155 observations were collected in the study, only 58 of them (∼37%) had data available for chlorophyte concentration. Finally, during part of the period of study, chlorophyte populations dropped to unusually low levels [22, 23], and outside of the periods of algal bloom (in all years), chlorophyte concentrations were often below the limit of detection, a problem which may be rectified with more modern equipment than that used in the study, which was conducted between 1994 and 2006. However, from an ecological standpoint, the algal bloom period is the most important time of year, since the algal bloom supports the GSL’s brine shrimp population. Therefore, moving forward, we will characterize the changes in water temperature and salinity during this time of year with priority, and measurements taken at other times of the year when the chlorophyte population is low may not be as important to consider. In terms of the analysis, both regression models that we developed have significant error, although the Poisson model is a better fit than the linear model. Fortunately, regression is a rich area of data analysis, and there are many more methods to try implementing to find the optimal fit. Furthermore, we may be missing variables in our analysis that help to explain the variability in the chlorophyte concentrations. By collecting additional data and fitting more regression models, we will find a more accurate prediction for *Dunaliella* concentrations moving forward.

### 4.2 Future Work

Future work on the GSL should accurately predict the South Arm’s microbial tipping point. The current models in this paper are not optimal and require refinement or refitting with another type of regression model in order to make more accurate predictions. Furthermore, predictions made in this work relied on observational data under conditions that are amenable to microbial growth. Experimental laboratory studies that probe extinction limits under co-varying conditions (e.g., elevated water temperature and salinity) should be conducted to collect a diverse array of data on *D. viridis*, such as growth rate, photosynthesis, and chlorophyll a.

Additionally, the gradual shift to higher salinity conditions in the North Arm since causeway construction (1959) could serve as a powerful precedent for the South Arm. It is currently unknown if observational data on phytoplankton populations (especially species) have been collected from the North Arm since 1959. If these data do exist, they will illuminate the shifts in microbial community structure with gradually increasing salinity and project potential impacts on the food web of the the South Arm.

Future work should also assess other abiotic factors impacted by climate change, such as increased mobilization of mercury in lake sediment and its brine layer. Greater bioavailability of mercury in the water column may decrease *Dunaliella* concentrations or increase the volume of individual cells, which may impact function at the population level [24].

### 4.3 Broader Implications

Saline lakes globally are a critical ecosystem that host species only found in these special environments. These ecosystems are also important economic drivers at the local, regional, and national level. However, saline lakes are disappearing at an alarming rate due to continued water diversion and climate change [11, 25]. Perhaps, by initiating a body of research to predict the collapse of *Dunaliella* populations and therefore the GSL ecosystem overall, this work will motivate a more immediate reaction from policymakers to preserve the Great Salt Lake.

## 5 Acknowledgements

This work was supported in part by the Interdisciplinary Quantitative Biology (IQ Biology) PhD program at the BioFrontiers Institute, University of Colorado Boulder, and the National Science Foundation NRT Integrated Data Science Fellowship (award 2022138).

We would like to thank Drs. Cresten Mansfeldt and Lizzy Trower for advising us on this project as well as Drs. Carie Frantz and Bonnie Baxter, who supplied their Great Salt Lake expertise, and Dr. Kristin Powell, Stephanie Hoyt, and the IQ Biology program at CU Boulder.

## 6 Supplementary Materials

For more information, visit [our GitHub page](https://github.com/rmb008/R4_GSLProject/tree/field_reg), which includes the dataset used in our analysis (field_data.csv) as well as the Python script used to conduct said analysis (regression.py).

